# Symbiont diversity within *Loripes orbiculatus* and the case for multiple hosts

**DOI:** 10.1101/2025.10.01.679794

**Authors:** Margaret A. Vogel, Fragkiskos Machairas, Jay Osvatic, Sophia Ferchiou, Bela Hausmann, Katja Klun, Jillian M. Petersen

**Affiliations:** Department of Fundamental Biology, University of Lausanne, Lausanne CH; Center for Microbiology and Environmental Systems Science, University of Vienna, Vienna AT; Joint Microbiome Facility of the Medical University of Vienna and the University of Vienna, University of Vienna, Vienna AT; Division of Clinical Microbiology, Department of Laboratory Medicine, Medical University of Vienna, Vienna AT; Marine Biology Station Piran, National Institute of Biology, Piran SL; Vienna Doctoral School in Microbiology and Environmental Science, University of Vienna, Vienna AT; Environment and Climate Hub, University of Vienna, Vienna AT

**Keywords:** Lucinid bivalves, Seagrass, Symbiosis, *Ca*.Thiodiazotropha, Facilitation

## Abstract

Seagrasses support immense biodiversity and are critical for maintaining coastal ecosystem health. These foundation species benefit from a ‘three-way’ facultative relationship with one of the common inhabitants of seagrass meadows, lucinid bivalves, which host specific bacterial *Ca*. Thiodiazotropha symbionts. Relatives of the bivalve symbionts have been detected on seagrass roots raising the possibility that these symbionts may colonize both animals and plants; however, no study has yet compared bivalve- and seagrass-associated symbionts at the same site and time. Our combination of 16S rRNA amplicon and metagenome sequencing revealed a greater diversity than was previously observed within both lucinid bivalves and on seagrass roots from the Adriatic Sea. We show that two of the *Ca*. Thiodiazotropha ASVs found on seagrass roots are identical to those found in bivalve hosts at the same site. This suggests that symbiont sharing may occur in the seagrass habitat between these two species, which has important evolutionary and ecological implications for both hosts and symbionts, as well as thin response to climate change.

## Introduction

The immense biodiversity found in seagrass meadows worldwide is supported in part by the dense habitat matrix created by seagrasses. These foundation species also provide valuable ecosystem services, including stabilizing sediments, oxygenating the water column, and blue carbon sequestration, making seagrasses critical for maintaining healthy coastal ecosystems. These highly productive systems also generate high rates of organic matter decomposition which can lead to increased and potentially phytotoxic levels of sulfide in the sediment [1]. Although seagrasses have evolved their own mechanisms to mitigate sulfide toxicity [2], they also rely on facilitative interactions with their inhabitants, including lucinid bivalves [3].

*Lucinidae* is a species-rich family of bivalves characterized by their symbiosis with sulfur-oxidizing Gammaproteobacteria that are housed within specialized cells called bacteriocytes in the gill tissue. These endosymbionts oxidize reduced sulfur and form a nutritional symbiosis with the bivalve host by delivering organic carbon. Lucinid bivalves have a global distribution and commonly occur within seagrass meadows [4,5]. Seagrasses and lucinids share a common evolutionary history having diversified concurrently in geological time [6]. These interactions persist today as their activity in seagrass beds is thought to facilitate seagrass growth. The chemical cycling performed by the symbionts in lucinid gills lowers sulfide levels in the sediment, which can, in turn, benefit the surrounding seagrass [4]. Not only has the presence of lucinids been shown to increase seagrass growth and survival under stressful conditions [3,7], but members of the genus *Candidatus* Thiodiazotropha, the group to which many lucinid symbionts belong, have also been found to be commonly associated with the roots of many seagrass species [8].

Although *Ca*. Thiodiazotropha has been studied separately in association with either marine plants or lucinid bivalves, no study has investigated its symbioses with both hosts where they co-occur (e.g., [5,9–12]). Here we report a fine-scale investigation of *Ca*. Thiodiazotropha diversity within 86 bivalve host individuals and several compartments of the surrounding seagrass environment at two sites in the Adriatic Sea. A combination of 16S rRNA amplicon and metagenome sequencing revealed a greater diversity than was previously observed within the bivalve gills as well as on seagrass rhizomes and roots. Additionally, this study is the first to show that select *Ca*. Thiodiazotropha ASVs found on seagrass roots and rhizomes are identical to those found in their bivalve host, *Loripes orbiculatus*, at the same site. This finding extends beyond previous research showing the relatedness of seagrass and bivalve symbionts by suggesting that some of these symbionts may be shared between these two hosts. Thus, seagrasses could act as a secondary host for this genus of symbionts, or that lucinid bivalves could act as secondary hosts for important symbionts of seagrasses.

## Methods

### Site Description

All samples were collected off the coast of Slovenia in the Gulf of Trieste at two nearby sites (Figure 1): Italian Border (IB; 45.592025°N, 13.715745°E) and Ankaran (AK; 45.572113°N, 13.741289°E). Both sites are characterized by seagrass habitats with *Cymodocea nodosa* (commonly known as little Neptune grass) as the dominant seagrass species. The Ankaran site also features a second distinct intertidal seagrass bed composed of *Zostera noltii* (commonly known as dwarf eelgrass) which becomes exposed at low tide.

**Figure 1.**
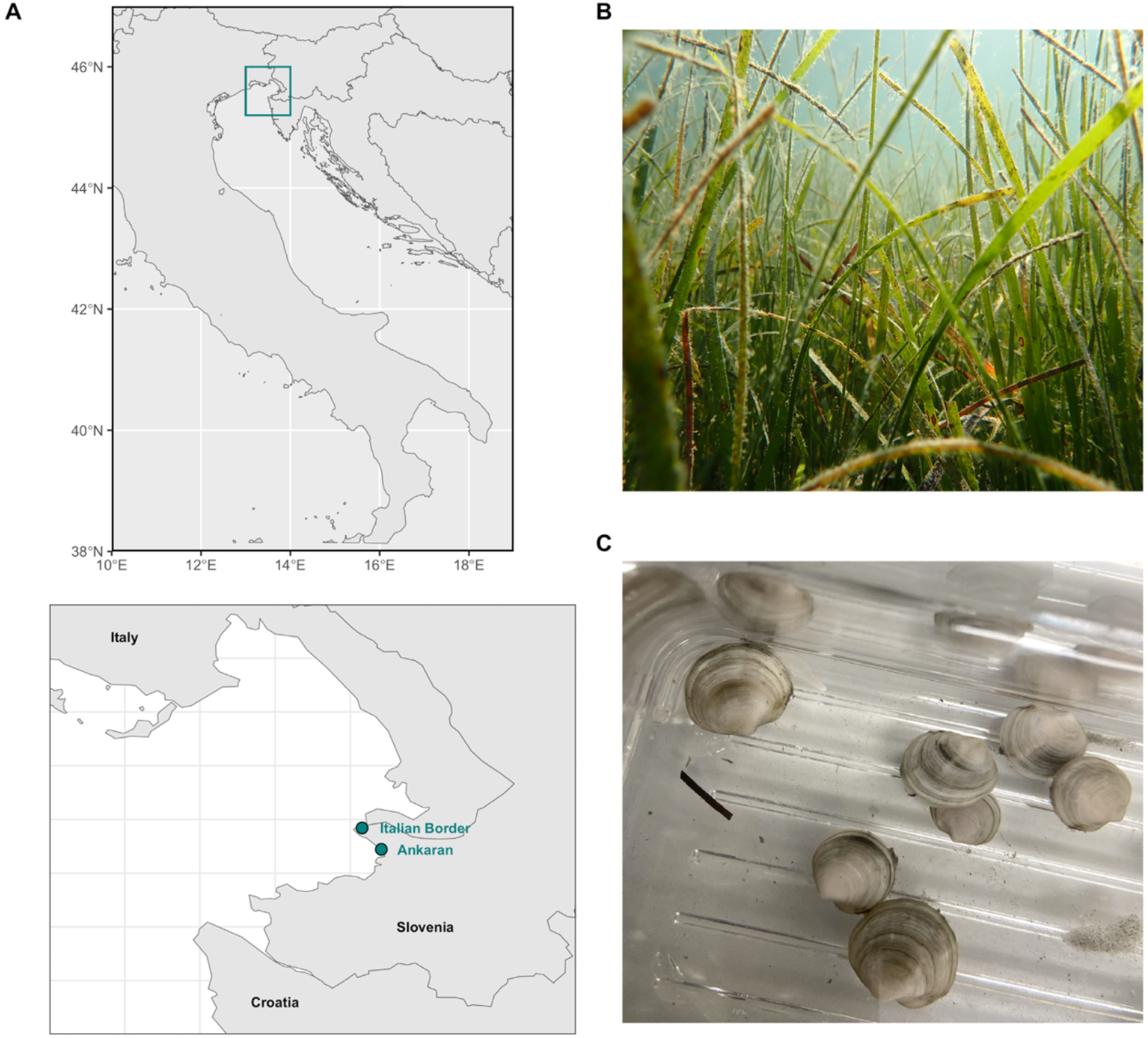
**(A)** Map of the two field sites, Italian Border and Ankaran, off the coast of Slovenia in the Adriatic Sea, **(B)** the submerged seagrass bed at Ankaran, and **(C)** *Loripes orbiculatus* taken from the field sites.

### Lucinid Sampling

To characterize the symbiont diversity at the two sites, *Loripes orbiculatus* individuals were collected from August to October 2021 from both sites (AK and IB). Sediment from in and around the seagrass beds at each site was collected into buckets and topped with seawater for transport back to the laboratory at the National Institute of Biology (NIB) Marine Biology Station Piran. Upon returning to the laboratory, sediments were sieved to isolate *L. orbiculatus*. The individuals collected for metagenome sequencing were also used in a separate experiment. All individuals were dissected and both gills were placed whole in RNAlater® and frozen at - 80°C.

### Seagrass Habitat Sampling

To identify environmental populations of *Ca*. Thiodiazotropha in the surrounding seagrass habitat, samples for microbial community profiling were collected at both sites from several compartments of the seagrass habitat: seagrass blades, seagrass roots and rhizomes, surrounding sediment, and overlying seawater. Two sampling events were conducted, one in July 2021 and the other in September 2021. During each sampling, five seagrass shoots were chosen haphazardly for sampling. One blade from each shoot was removed using sterile forceps and a sterile swab was used to sample the microbial communities following the methods of Vogel et al. [13]. Pieces of rhizome and the attached roots were collected from five separately chosen shoots. The root and rhizome fragments were detached using sterile forceps and placed into sterile tubes. All blade and root/rhizome samples from the Italian Border site were collected from *C. nodosa*, while at the Ankaran site, five of each sample type was collected from *C. nodosa* shoots and five from *Z. noltii*. Additionally, at each site, sediment samples were collected near seagrass roots and placed into sterile tubes. Water column microbial communities were sampled from seawater overlying the seagrass bed using a sterile syringe and microbial biomass was collected on a 0.22 μM Sterivex ™ filter.

Seagrass blades, seagrass roots and rhizomes, and seawater were immediately fixed in in RNAlater® and all samples were placed on ice to be transported to the laboratory at the National Institute of Biology (NIB) Marine Biology Station Piran. Samples were kept at -80°C before transport in a dry shipper cooled with liquid nitrogen to the laboratories at the University of Vienna where they were again stored at -80°C until further processing.

During the September 2021 sampling, additional samples were collected to obtain seawater and porewater nutrient concentrations at both sites. For seawater nutrient analysis, samples were collected from the seawater overlying the seagrass bed at each site and filtered through a 0.22 μM filter. Sediment cores were collected at each site and porewater was extracted from the total length of the core. The collected water samples for nutrient analysis were stored in acid-washed 15 mL tubes and immediately frozen at -20°C. Nutrient analysis (nitrate, nitrite, ammonia, and phosphate) was conducted at the NIB Marine Biology Piran Station using a segmented flow analyzer (QuAAtro, SealAnalytical, [14]). Quality control of analysis was done with certified reference material (CRM) for nutrients in seawater (KANSO TECHNOS) and by annual participation in intercalibration program (WEPAL-QUASIMEME).

### Microbial Community Analysis

Genomic DNA (gDNA) was extracted from all environmental samples (seagrass blades, seagrass roots and rhizomes, sediment, and seawater) using the DNeasy PowerSoil Pro Kit (Qiagen, Hilden, Germany) following the manufacturer’s instructions. The gill tissue for each *L. orbiculatus* individual was separately homogenized and gDNA was extracted using the All-In-One DNA/RNA/Protein Miniprep Kit (Bio Basic, Ontario, Canada). All gDNA extracts were transferred to the Joint Microbiome Facility (JMF) of the Medical University of Vienna and University of Vienna where 16S rRNA gene iTag libraries were prepared for all samples using the archaeal and bacterial primers 515F and 806R (targets the V4 region of *E. coli*) modified by Apprill et al. [15] and Parada et al. [16]. Amplicons were then sequenced using an Illumina MiSeq and the raw sequences were processed with DADA2 following the protocols described in Pjevac et al. [17]. The constructed amplicon sequence variant (ASV) table was then filtered to remove any sequences resulting from mitochondrial, chloroplast, or eukaryotic DNA. Taxonomy was assigned using the SINA classifier with SILVA reference database v.138. Samples with poor sequence quality, including 13 root/rhizome samples and one gill sample, were excluded from further analysis. The resulting ASV table was then normalized using cumulative sum scaling (CSS) to account for differences in sequencing depth among samples. All statistical analyses were conducted on the CSS normalized ASV table using R v. 4.2.3. For a subset of the gill samples (86/130), metagenomic sequencing was also performed using short-read sequencing using an Illumina MiSeq. Individual read libraries were quality checked using fastQC v. 0.12.1. Adapters were trimmed and phiX contamination was removed using BBDuk (part of BBMap v. 39.10) to a minimum of 50 base pairs in length. Reads were k-trimmed from the right with a kmer of 21, minimum kmer of 11 and hamming distance of two along with the “tpe” and “tbo” options. Quality trimming was performed from the right with a Q-score of 15. BBMap’s reformat.sh was used to interleave read libraries and libraries were merged. Phyloflash v. 3.4.2 was used to generate 16 and 18 SSU rRNA sequences using the SIVLA v. 138.1 database. The interleaved library was assembled using SPAdes v. 4.1.0 in “meta” mode and BBMap’s reformat.sh was used to remove contigs under 1000 base pairs from assembly. Prodigal v 2.6.2 in “meta” mode was used to extract all genes from the assemblies.

Metagenomic reads were aligned to a custom reference database comprising complete mitochondrial genomes and *cytochrome b* gene sequences from Lucinidae species using Bowtie2 [18] to identify candidate host mitochondrial sequences. Matching reads were deduplicated using fastp [19] and subsequently assembled *de novo* into contigs using SPAdes [20] in single-end mode. Contigs shorter than 1,000 base pairs were excluded from further analysis. The remaining contigs were remapped to the reference database using minimap2 [21], and the aligned regions were extracted using bedtools [22]. For each sample, the mapped regions were concatenated into a single representative mitochondrial sequence. All sequences were aligned using MAFFT [23], and phylogenetic trees were inferred using FastTree [24].

## Results

### *Loripes orbiculatus* gill microbial communities

Microbial communities from the homogenized gill tissues of 130 lucinid individuals contained a total of 164 ASVs and had an average richness of 31.8 ASVs. These communities were mainly comprised of *Ca*. Thiodiazotropha spp., the known symbionts of *L. orbiculatus*, with 20 ASVs belonging to this genus accounting for 95% of the relative abundance on average in all individuals. The number of *Ca*. Thiodiazotropha ASVs in each lucinid bivalve ranged from 4-13 with a mean of 7.7 ASVs observed. Additionally, there were three *Ca*. Thiodiazotropha ASVs (ASV_7xh_i9e, ASV_imh_tqk, and ASV_ey3_jh5) made up most of this fraction, with at least one of these ASVs dominating in each lucinid individual (Figure 2A). Both the sequence from ASV_7xh_i9e and from ASV_imh_tqk had a 100% match to the 16S rRNA region of a species (*Ca*. Thiodiazotropha lotti and *Ca*. Thiodiazotropha luna, respectively) from metagenome-assembled genomes (MAGs) previously sampled from lucinid gills [5,12]. The third dominant ASV (ASV_ey3_jhf) also had a very high sequence similarity to *Ca*. T. luna and ASV_imh_tqk with a single base pair difference between the two ASVs.

**Figure 2.**
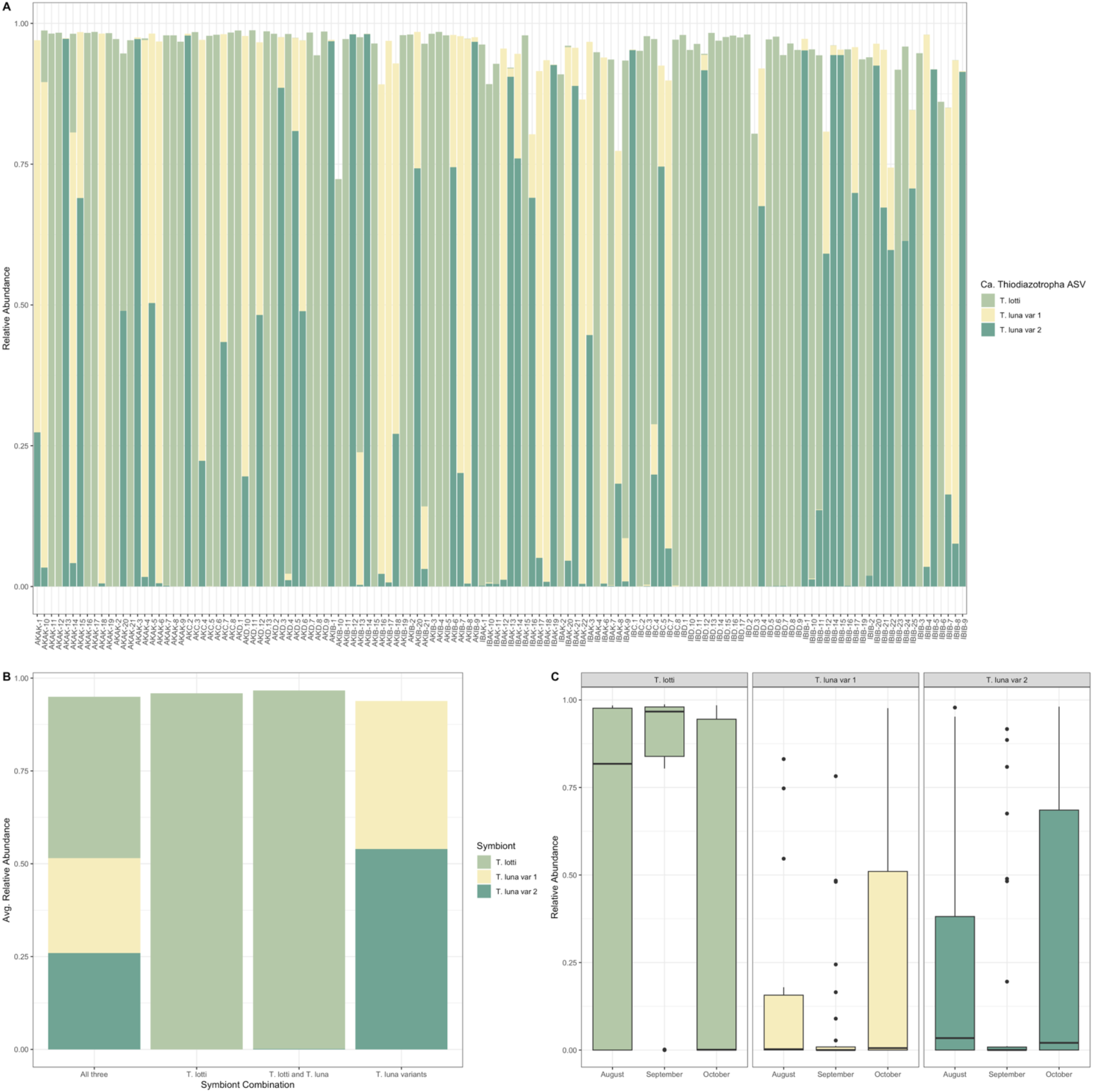
**(A)** Relative abundances of the three dominant symbiont sequence types in all lucinids (n=130). Colors indicate the symbiont sequence type that has the largest proportional abundance in each sample. Delete x-axis labels, **(B)** co-occurrence patterns of the three symbiont sequence types. Add in occurrence percentage, and **(C)** relative abundances of the three dominant symbiont sequence types by sampling date.

Metagenomes were available from individuals where one of these two very similar ASVs dominated 16S rRNA amplicon libraries, allowing their genomes to be compared based on the MAGs assembled from each metagenome. These MAGs showed only 96.5% ANI similarity between these two symbiont sequence types. Therefore, these ASVs likely represent ‘variants’ of one another and will be referred to as *Ca*. T. luna variant 1 (ASV_imh_tqk) and *Ca*. T. luna variant 2 (ASV_ey3_jhf). MAG comparisons also confirmed the identities of *Ca*. T. lotti in samples and showed a distinct difference between *Ca*. T. lotti and both *Ca*. T. luna variants with a 78.3-79.9% ANI similarity.

We investigated the relationship between host mitochondrial variation and symbiont composition using both complete mitochondrial genomes and the *cytochrome b* gene region. After excluding samples with assemblies shorter than 1,000 bp, 34 high-quality samples remained for phylogenetic analysis. Across these samples, all mitochondrial and *cytochrome b* sequences were consistent with those of *Loripes orbiculatus*, confirming the species identity. To test whether host genotype (i.e., mitochondrial phylogeny) was associated with symbiont community structure, we examined both the dominant symbiont taxon and overall symbiont diversity. Mantel tests showed no significant correlations between host phylogenetic distances and either symbiont richness (r ≈ 0, p > 0.05) or relative abundance profiles. Similarly, Pagel’s λ tests for phylogenetic signal indicated no significant association between host phylogeny and the distribution of dominant *Ca*. Thiodiazotropha ASV (λ ≈ 0, p > 0.05), suggesting that host genotype does not constrain symbiont identity or diversity in this system.

In the 130 *Loripes orbiculatus* individuals sampled, *Ca*. T. lotti was dominant in 67 individuals, while *Ca*. T. luna variant 2 was dominant in 35 and *Ca*. T. luna variant 1 dominant in 28 individuals. The three dominant symbiont sequence types exhibited complex co-occurrence patterns indicating distinct interactions between them (Figure 2A). *Ca*. T. lotti occurred alone in 23.8% of lucinid hosts and is the only one of the three dominant symbiont sequence types that occurred alone without a detectable amount of one of the two others. In a small proportion of individuals (14.6%), *Ca*. T. lotti occurred at a low relative abundance with one of the *Ca*. T. luna variants and in around 25% of the host individuals, all three dominant symbiont sequence types occurred. In the remaining 36.9% of hosts, the *Ca*. T. luna variants occurred with one another in varying proportions, however, neither of the two variants were ever detected without the other (Figure 2B). The dominant *Ca*. Thiodiazotropha ASVs comprised an average of 87.9% ± 1.18% SE (13.4% SD) of the total microbial relative abundance in each bivalve. *Ca*. T. lotti ranged from 68.3% to 98.7% relative abundance, while the relative abundance of the *Ca*. T. luna variants ranged from 48.4% to 98.1%. The lower range of *Ca*. T. luna variants can be explained by their occurrence always together.

Shell size of bivalves collected in October (n=86) was measured and was significantly different by site with individuals from Ankaran having a larger shell size on average (11.74 ± 2.32 mm, SD) than individuals from the Italian Border (6.15 ± 0.65 mm, SD; Kruskal-Wallis, χ^2^=62.945, df=1, p<0.001). However, the dominant symbiont sequence type nor the relative abundance of the dominant symbionts did not significantly differ with shell size (Kruskal-Wallis, p>0.05). Further, in lucinids collected from all sampling dates, no significant differences in dominant symbiont sequence type or the relative abundance of the three symbiont sequence types between sites were detected (Kruskal-Wallis, p>0.05, Figure 2C). However, there was a significant difference in the dominant symbiont sequence type among sampling dates with significantly greater relative abundances of *Ca*. T. lotti observed in individuals collected in September than in October (Kruskal-Wallis, χ^2^=15.747, df=2, p<0.001; Dunn’s test with Benjamini-Hochberg adjusted p<0.001). Conversely, greater relative abundances of *Ca*. T. luna variant 1 and *Ca*. T. luna variant 2 (Kruskal-Wallis, p<0.05; Dunn’s test with Benjamini-Hochberg adjusted p<0.05; Figure 2C) were observed in individuals collected in October. These changes in the relative abundance of the dominant symbiont sequence types over time could be driven by seasonal changes in environmental conditions; however, this dataset is limited by environmental parameters measured only in September.

In September, the Italian Border site had significantly higher concentrations of nitrate (Kruskal-Wallis, χ^2^=7.0313, df=1, p=0.008) and ammonium in the overlying seawater (Kruskal-Wallis, χ^2^=6.8598, df=1, p=0.008) as well as significantly higher sediment porewater concentrations of ammonium (Kruskal-Wallis, χ^2^=4.5, df=1, p=0.034) and phosphate (Kruskal-Wallis, χ^2^=4.6667, df=1, p=0.031; Table 1; Figure S1). During the September sampling, *Ca*. T. lotti was dominant in more bivalve individuals from the Italian Border (88%) compared to individuals from Ankaran (62%). Additionally, at the Italian Border, *Ca*. T. luna variant 1 was never the dominant symbiont sequence type; yet at Ankaran, *Ca*. T. luna variant 1 was dominant in 15% of bivalves. However, the dominant symbiont sequence type and relative abundances of the symbiont sequence types did not significantly differ by site in September (Kruskal-Wallis, p>0.05). This suggests that the site-specific variables measured here, including nutrient concentrations or seagrass species present, are not major determinants of which *Ca*. Thiodiazotropha symbiont sequence type dominates the symbiont community in individual lucinids.

**Table 1.**
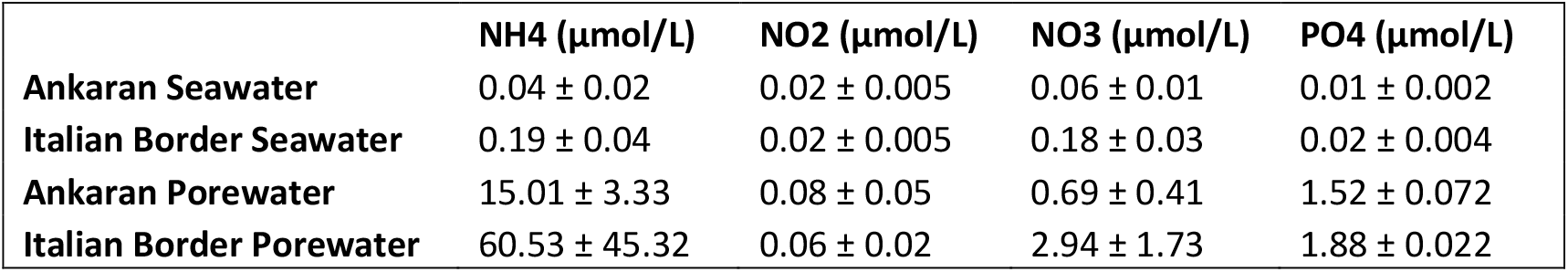
Water column and sediment porewater nutrient concentrations at the Ankaran and Italian Border sites. Values are mean ± standard deviation.

The non-dominant remaining proportion of *Ca*. Thiodiazotropha ASVs comprised 0.17% to 18.4% in combined relative abundance. As expected, compositional differences in the entire gill microbial community were best explained by the dominant symbiont sequence type due to their high relative abundance and low community evenness (PERMANOVA, R^2^=0.76477, F=206.45, p=0.001; Figure S2). Interestingly however, when the dominant symbiont sequence types were omitted from the analysis, the remaining gill community still showed significantly different composition due to the dominant symbiont sequence type and marginally significant differences due to site with no interaction between the two (PERMANOVA, p=0.001, p=0.086; see Table 2) and distinct clustering of those communities dominated by *Ca*. T. lotti and dominated by the two *Ca*. T. luna variants remained (Figure 3). This suggests that the dominant symbiont sequence type may be interacting with the bivalve host or modifying the gill microenvironment to shape the overall gill microbial community.

**Table 2.** PERMANOVA results for the remaining gill microbial communities without the three dominant symbiont types (*Ca*. T. lotti, *Ca*. T. luna var 1, *Ca*. T. luna var 2) showing composition of communities significantly differs by the dominant symbiont sequence type and, to a lesser degree, site.

|  | Df | Sum of Sqs | R2 | F | Pr(>F) |
| --- | --- | --- | --- | --- | --- |
| Site | 1 | 0.3167 | 0.01011 | 1.7297 | 0.086 |
| Symbiont Sequence Type | 2 | 7.9232 | 0.25304 | 21.6355 | 0.001*** |
| Site* Symbiont Sequence Type | 2 | 0.3667 | 0.01171 | 1.0013 | 0.440 |
| Residual | 124 | 22.7053 | 0.72513 |  |  |
| Total | 129 | 31.3120 | 1 |  |  |

**Figure 3.**
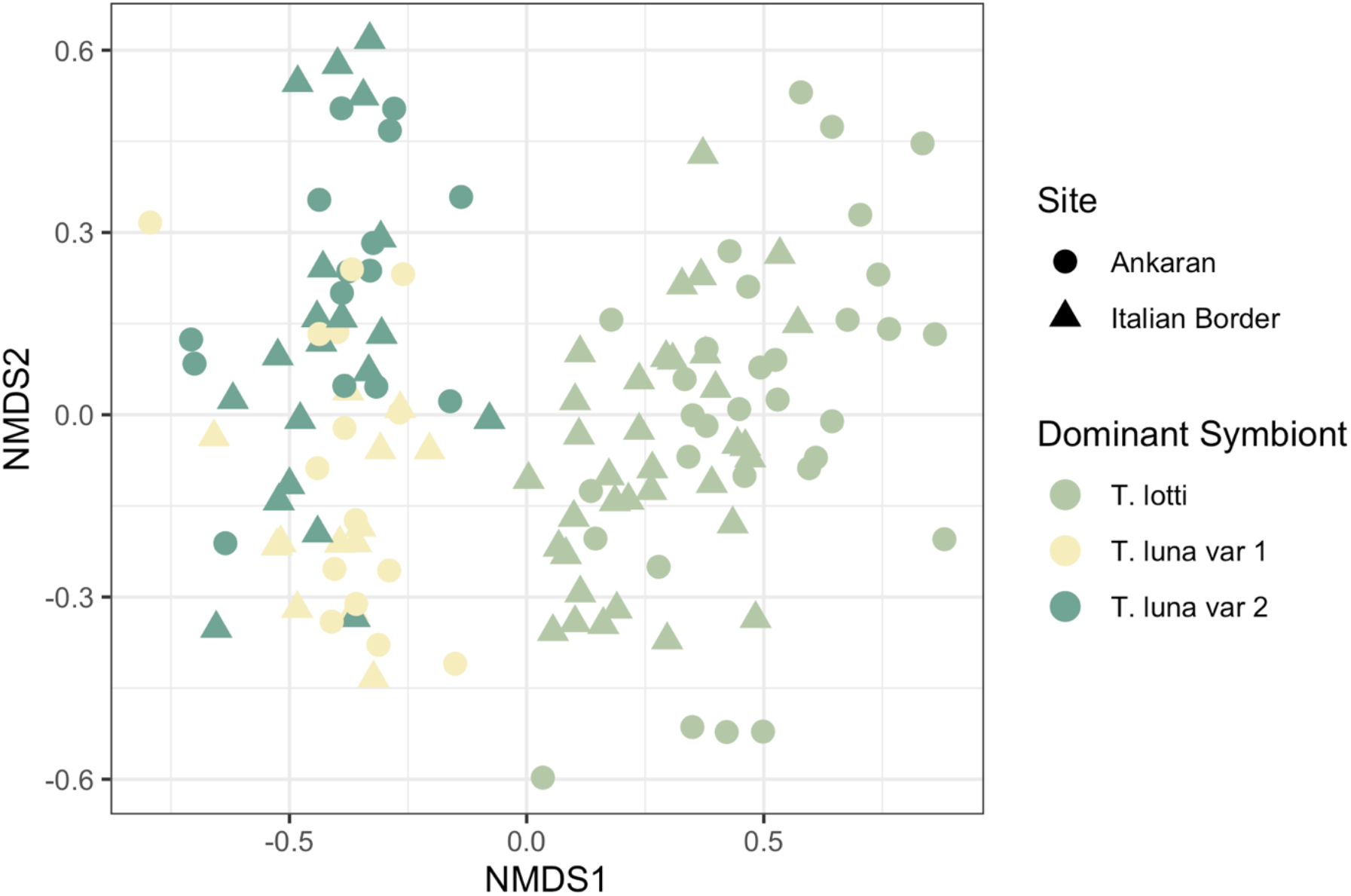
Non-metric Multi-Dimensional Scaling ordination of the gill microbial communities without the three dominant symbiont sequence types (*Ca*. T. lotti, *Ca*. T. luna var 1, *Ca*. T. luna var 2) showing composition is significantly different by dominant symbiont sequence type even when they are excluded from the species abundance matrix (Bray-Curtis distance, k=3, stress=0.1695798).

### *Ca*. Thiodiazotropha spp. in the surrounding seagrass habitat

Microbial communities from the different compartments of the seagrass habitat (seagrass blade, seagrass root and rhizome, seawater and sediments) had a distinct composition and structure (PERMANOVA, R^2^=0.46605, p=0.001; Figure 4). Although ASVs belonging to *Ca*. Thiodiazotropha were found in all compartments of the environment at low abundances, the root and rhizome had significantly higher relative abundances of *Ca*. Thiodiazotropha ASVs than all other sample types (Kruskal-Wallis, χ^2^=31.689, df=3, p<0.001; Dunn’s test with Benjamini-Hochberg adjusted p<0.05; Figure 5A). The microbial communities on the roots and rhizomes also had the highest *Ca*. Thiodiazotropha richness with 11 of the 13 ASVs found in the seagrass environment observed on the roots and rhizomes (Figure 5B). The higher richness and abundances of *Ca*. Thiodiazotropha ASVs observed on the roots and rhizomes is consistent with previous findings that this genus of sulfur-oxidizing bacteria are consistent members and enriched in the seagrass rhizosphere and possibly engage in a beneficial symbiosis with the seagrass host [8]. Two of the *Ca*. Thiodiazotropha ASVs (*Ca*. T. luna variant 1 and ASV_1n2_n6n) found in the seagrass habitat were also detected in the gill tissue (Figure 6). *Ca*. T. luna variant 1 was detected in all compartments of the seagrass habitat while ASV_1n2_n6n was only detected on the seagrass blades and rhizome. ASV_1n2_n6n occurred in ∼60% of all lucinid individuals and was found at a slightly higher abundance in the rhizome (∼0.04%) than *Ca*. T. luna variant 1 (∼0.02%), although *Ca*. T. luna variant 1 occurred more frequently in a greater number of samples. Additionally, many of the other *Ca*. Thiodiazotropha ASVs that were not observed in the lucinid gill, were more prevalent on the seagrass root/rhizome than the two shared ASVs (Figure 5A). Yet, this shows that *Ca*. T. luna variant 1 and ASV_1n2_n6n can utilize both the seagrass and lucinid bivalves as a host and suggests symbiont sharing may be occurring within the seagrass habitats.

**Figure 4.**
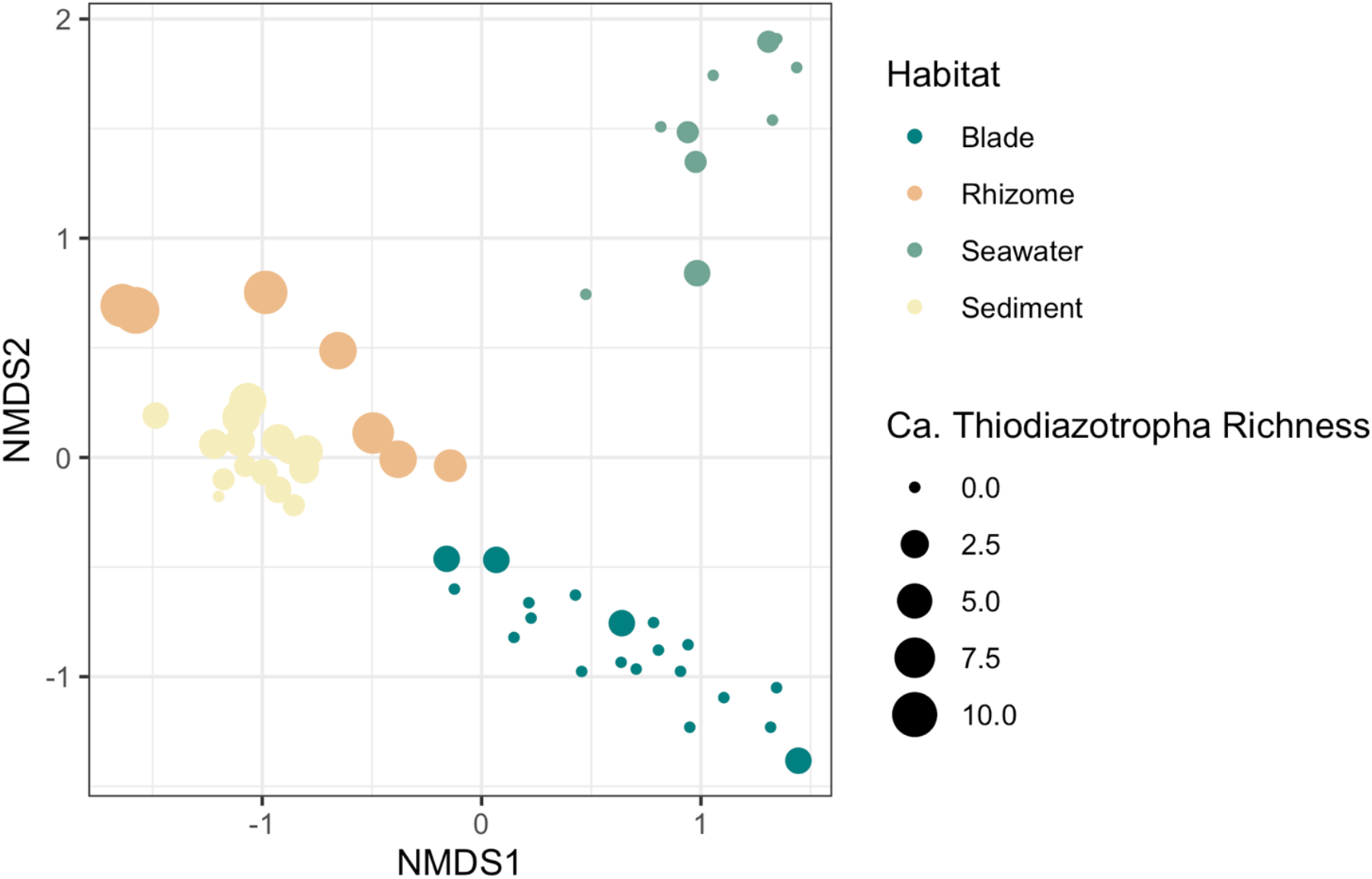
Non-metric Multi-Dimensional Scaling ordination of microbial communities from the seagrass habitat. Colors indicate the habitat compartment and size of the point indicates the observed richness of *Ca*. Thiodiazotropha ASVs in the community (Bray-Curtis distance, k=3, stress=0.06764439).

**Figure 5.**
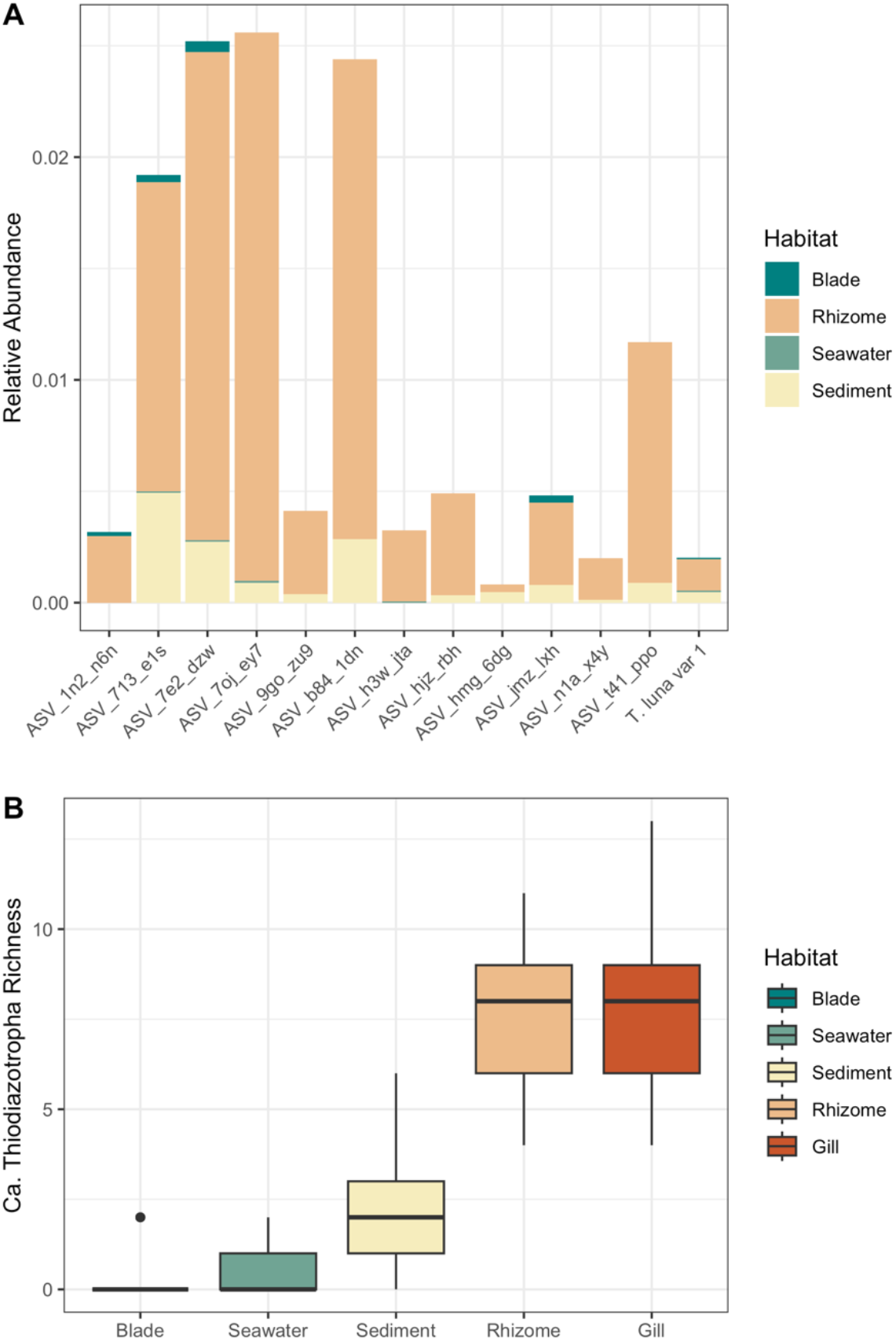
**(A)** Relative abundance of *Ca*. Thiodiazotropha ASVs in the four compartments of the seagrass habitat showing significantly greater abundance in the seagrass rhizome and **(B)** richness of *Ca*. Thiodiazotropha ASVs in the seagrass habitat and lucinid gills.

**Figure 6.**
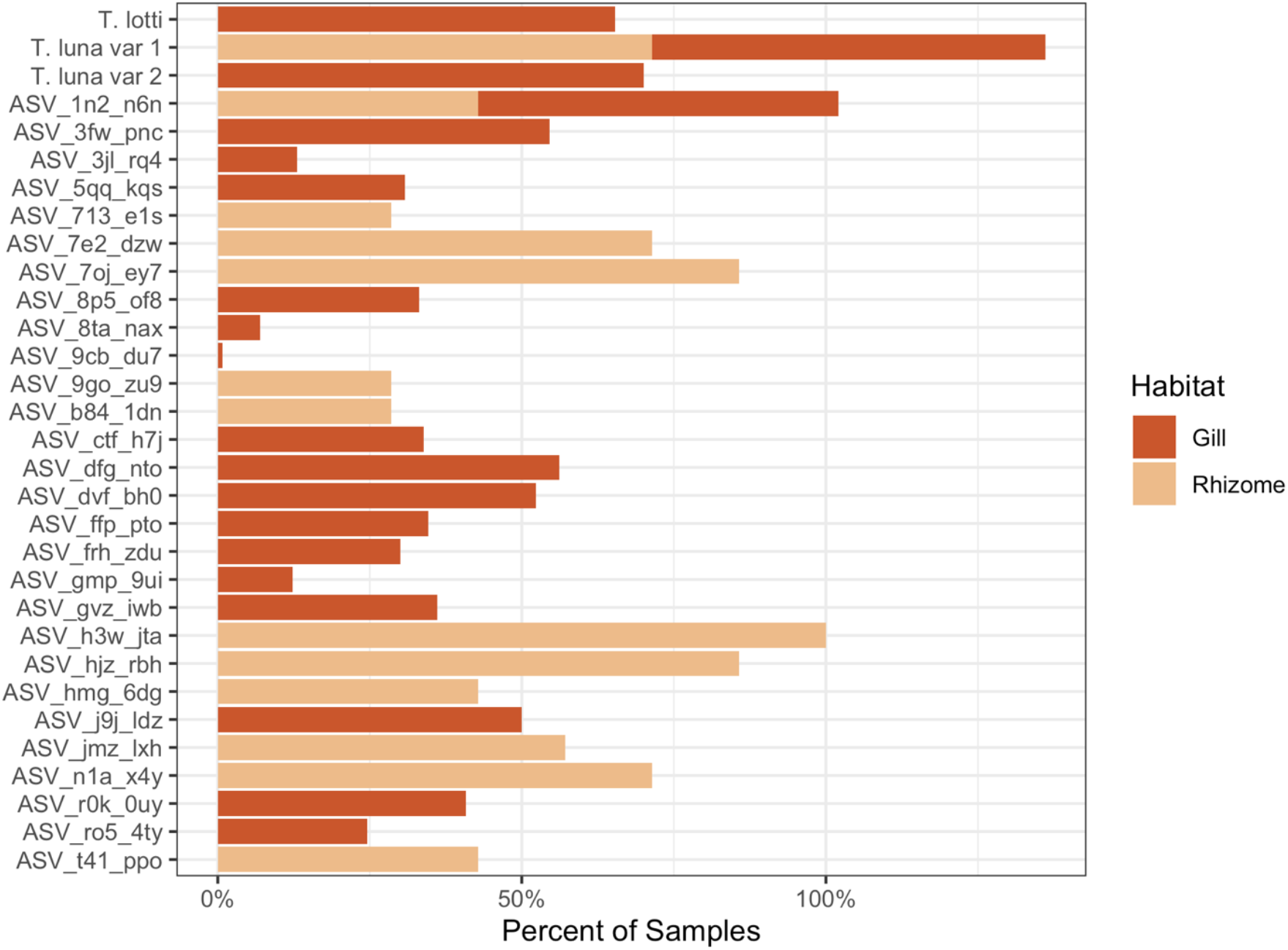
Occurrence of all *Ca*. Thiodiazotropha ASVs observed in the gill and rhizome with *Ca*. T. luna variant 1 and ASV_1n2_n6n occurring in both habitat types.

## Discussion and Conclusion

This study showed that a greater *Ca*. Thiodiazotropha diversity occurs in *Loripes orbiculatus* than previously observed with three dominant symbiont sequence types commonly co-occurring in the population and within a single individual. Further, a new symbiont sequence type, *Ca*. T. luna variant 2, was identified and was observed to occur frequently and sometimes at high abundance in individual bivalves at the two sites. Lim et al. [9] observed similar fine-scale diversity patterns in the lucinid bivalve *Ctena orbiculata* with four coexisting *Ca*. Thiodiazotropha-like symbiont sequence types in the population. The observed abundances of the three dominant *Ca*. Thiodiazotropha symbionts in this study did not differ with respect to the two sites despite the sites having differences in porewater and water column nutrient concentrations, shell size, and seagrass species present at the site. The dominant symbiont sequence type differed by sampling date, which could be driven by environmental variables not measured or by changes to gill morphology due to the reproductive cycle. In *Loripes orbiculatus*, the ratio of bacteriocytes to mucocytes can decrease during warm periods, which could affect patterns of symbiont relative abundance, and is thought to coincide with spawning periods [25].

Results also show that the dominant symbiont sequence type present in a lucinid individual may structure the remaining bacterial community on and within the gill. This suggests that the dominant symbiont sequence types may be interacting differently with their bivalve host, altering its immune responses, modifying the gill microenvironment, or directly interacting with co-occurring microbes in the gill. In the future, experimental data such as metatranscriptomes from *L. orbiculatus* hosting different symbiont sequence types could shed light on their possible functional differences.

As *Ca*. Thiodiazotropha symbionts are horizontally transferred from parent to offspring through the environment by *Loripes orbiculatus* and other lucinid species [26–28], it has been long assumed that these symbionts must have an environmental reservoir. Many ASVs belonging to *Ca*. Thiodiazotropha were observed in the seagrass habitat with significantly higher abundances on the seagrass roots and rhizomes than in the other environmental compartments sampled. These results confirm a previous meta-analysis of seagrasses worldwide [8] and suggest that *Ca*. Thiodiazotropha may be specific and permanent members of the seagrass rhizosphere microbial community. This association could enable seagrasses to mitigate harmful levels of sulfide in sediments with no lucinid bivalves present, and may be critical for optimal seagrass growth, particularly considering that some *Ca*. Thiodiazotropha are able to fix nitrogen, often limiting for seagrass growth [29–32]. A similar symbiosis has been demonstrated in the salt marsh grass *Spartina alterniflora* where close relatives of *Ca*. Thiodiazotropha contribute to sulfur oxidation on the roots with increased levels of expression of sulfur-oxidizing genes (oxidative-dsrAB) on roots under sulfide stress [33]. The extremely close relationship between bivalve- and seagrass-associated *Ca*. Thiodiazotropha, with identical 16S rRNA genes in the region we sequenced, shows that symbiont sharing between hosts within the seagrass habitat is occurring, even if additional genomic data would be needed to show whether truly genetically identical symbionts can colonize both hosts, or if minor genetic changes are required to adapt to association with different hosts, as was previously shown with a single promoter in *Vibrio fisheri* colonizing squid or fish [34].

Lucinid bivalves, their symbionts, and numerous seagrass species are involved in a three-way facilitative relationship resulting from millions of years of shared geologic and evolutionary history. Yet, it is unclear how rapid changes in the environment conditions due to climate change will affect these facilitative interactions and coevolution [35]. A previous study showed that the genetic diversity of the chemosynthetic endosymbionts associated with a genus of marine snails (*Alviniconcha*) was due to adaptation of the symbionts to local environmental conditions rather than host genetics [36]. Association of the host organism with locally adapted symbionts may provide the host with fitness advantages, especially under potentially stressful conditions. In this way, the association of both bivalves and seagrasses with *Ca*. Thiodiazotropha symbionts may allow these hosts to better mitigate degraded environmental conditions and these facilitative interactions may persist or even strengthen under climate change.

However, it is also possible that the effects of climate change will cause a weakening in these interactions. Seagrass coverage is already declining at an unprecedented rate due to anthropogenic activities, and this loss is often coupled with a decrease in seagrass diversity as species composition shifts from long-lived, climax species to shorter-lived, opportunistic species [37,38]. The decrease in abundance and loss of biodiversity of the seagrass host could reduce the amount of symbiont sharing within the seagrass habitat, thus weakening seagrass-symbiont interactions. Further, the reduced availability of a second host could result in the loss of genetic variation within the symbiont populations and, in turn, affect the interactions between *Ca*. Thiodiazotropha and the bivalve host. The amount and directionality of symbiont sharing needs to be determined in order to assess how changes in host availability could affect symbiont diversity and their relationships to both hosts to better predict the long-term effects of climate change [39–41].

We showed that two of the *Ca*. Thiodiazotropha ASVs found on the seagrass roots and rhizomes were also found in the gill tissue of *Loripes orbiculatus* at the same site and time. This reveals not only that seagrass roots may offer a suitable niche for *Ca*. Thiodiazotropha as well as certain species of *Ca*. Thiodiazotropha may be able to use both seagrass and bivalves as hosts. Symbiont sharing between two different hosts in a shared habitat has important implications for *Ca*. Thiodiazotropha ecology and evolution and for the interactions between the two hosts. For example, it may support transmission between host generations or maintain a higher diversity of symbionts in both the bivalve and the seagrass hosts. This diversity, in turn, may enable seagrasses to endure a broader range of stressful conditions and promote growth, underpinning its essential ecosystem services. To build upon this, future studies should investigate the functions *Ca*. Thiodiazotropha performs on seagrass roots and rhizomes and how they interact with the seagrass host. A better understanding of how these symbionts move through the environment is also needed to determine the time scale of host switching in these systems and how such closely related or possibly even identical organisms manage to successfully colonize the intracellular environment in animals as well as the surfaces of plant roots. Such intimate and beneficial associates as *Ca*. Thiodiazotropha that colonize host animal gill cells are usually highly specific for their co-evolved host organism (e.g., [42]). Although some bacterial pathogens can associate with hosts across an equally broad taxonomic range (e.g., [43,44]), this would be an unprecedented ability for beneficial symbionts.

## Acknowledgments

The work was supported by a WWTF Vienna Research Grant, an ERC Starting Grant EvoLucin and ERC Consolidator Grant SeaSym, as well as the Austrian Science Fund (FWF; 10.55776/COE7; the MAINTAIN Doc.Funds project). M.A.V. was additionally supported by Swiss National Science Foundation Eccellenza grant PCEGP3_181272 and the Swiss National Science Foundation National Center of Competence in Research Microbiomes grant 51NF40_180575. Funded by the European Union. Views and opinions expressed are however those of the author(s) only and do not necessarily reflect those of the European Union or the European Research Council Executive Agency. Neither the European Union nor the granting authority can be held responsible for them.

## Data Availability

Submission of all associated data to public databases, including NCBI, is in progress. All data will be made publicly available before acceptance and can be provided at any time to reviewers if requested.

## Supplementary Figures

**Figure S1.**
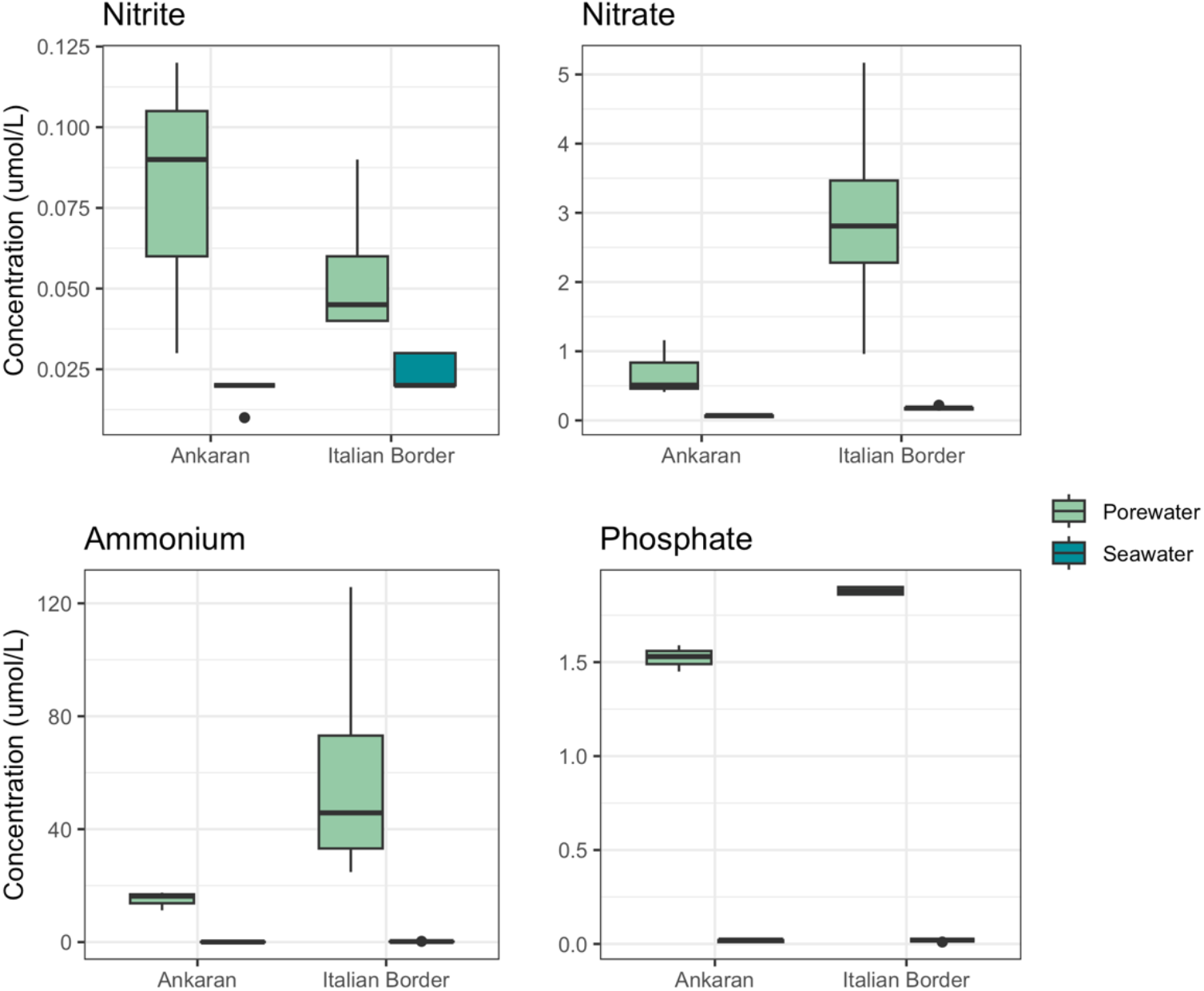
Nutrient concentrations (nitrate, nitrite, ammonium, and phosphate) in the porewater and overlying seawater at both sites.

